# Genome-Wide Analysis of Single Non-Templated Nucleotides in Plant Endogenous siRNAs and miRNAs

**DOI:** 10.1101/044982

**Authors:** Feng Wang, Nathan R. Johnson, Ceyda Coruh, Michael J. Axtell

**Affiliations:** Intercollege Plant Biology Ph.D. Program, Huck Institutes of the Life Sciences, Penn State University, University Park, PA 16802 USA.; Department of Biology, Penn State University, University Park, PA 16802 USA

**Author notes:** Current address: Salk Institute for Biological Studies, La Jolla, CA 92037 USA.

## Abstract

Plant small RNAs are subject to various modifications. Previous reports revealed widespread 3' modifications (truncations and non-templated tailing) of plant miRNAs when the 2'-C-methyltransferase HEN1 is absent. However, non-templated nucleotides in plant heterochromatic siRNAs have not been deeply studied, especially in wild-type plants. We systematically studied non-templated nucleotide patterns in plant small RNAs by analyzing small RNA sequencing libraries from *Arabidopsis*. tomato, *Medicago*, rice, maize, and *Physcomitrella*. Elevated rates of non-templated nucleotides were observed at the 3' ends of both miRNAs and endogenous siRNAs from wild-type specimens of all species. 'Off-sized' small RNAs, such as 25 and 23 nt siRNAs arising from loci dominated by 24 nt siRNAs, often had very high rates of 3'-non-templated nucleotides. The same pattern was observed in all species that we studied. Further analysis of 24 nt siRNA clusters in *Arabidopsis* revealed distinct patterns of 3'-non-templated nucleotides of 23 nt siRNAs arising from heterochromatic siRNA loci. This pattern of non-templated 3' nucleotides on 23 nt siRNAs is not affected by loss of known small RNA 3'-end modifying enzymes, and may result from modifications added to longer heterochromatic siRNA precursors.

## Introduction

Plant regulatory small RNAs, which are usually 20 - 24 nt in length, are classified into different sub-categories based on their biogenesis and function (1). Small RNA genes can be empirically annotated by the pattern of alignments and size of predominant small RNAs (2, 3). Plant microRNAs (miRNAs), usually 21 nt in length, and heterochromatic small interfering RNAs (het-siRNAs), usually 24 nt in length, are the two major types of plant small RNAs. Plant miRNAs are processed from single stranded hairpin-forming primary RNAs and mediate post-transcriptional silencing by triggering target mRNA slicing and/or translational repression (4). Plant het-siRNAs are processed from double stranded RNA and mediate silencing by RNA-directed DNA methylation (RdDM) (5).

Though plant miRNAs and het-siRNAs have different biogenesis and function pathways, they both confer silencing through base-pairing interactions with target RNAs (6-8). Posttranscriptional modifications of small RNAs can lead to reduced targeting specificity and/or altered small RNA stability (9). In plants, both miRNAs and het-siRNAs are subject to 2'-*O*-methylation of the 3'-most ribose by the small RNA methyltransferase HUA ENHANCER 1 (HEN1), which protects plant small RNAs from degradation (10, 11). Small RNAs in *hen1* mutant backgrounds are subject to extensive 3' to 5' exonucleolytic truncation and/or 3' tailing (4). Various 3' end nucleotide addition patterns of plant miRNAs have been observed in *hen1* mutants as well as in wild-type plants. 3' end uridylation of unmethylated miRNAs, which is the predominant 3' tailing form in *Arabidopsis hen1* plants, usually signals destabilization of miRNAs. HEN1 Repressor 1 (HESO1) is the primary uridyltransferase for uridine addition on the 3' ends of miRNAs (12, 13). UTP:RNA Uridyltransferase 1 (URT1) cooperatively uridylates miRNAs along with HESO1 in *Arabidopsis* (14, 15). Other than triggering miRNA destabilization, uridylation also can affect the function of miRNAs. A uridylated 22 nt variant of miR171a that only arises in a *hen1* mutant background can trigger secondary phased siRNA biogenesis (14, 16). Non-templated nucleotides other than U are also observed in the *hen1* background, but their biological functions are less well understood. Interestingly, Lu *et al*. observed that 3' adenylated miRNAs in *Populus trichocarpa* seemed to show a slower degradation rate (17).

Much less is known about the patterns of non-templated nucleotides of het-siRNAs. Like miRNAs, het-siRNAs are HEN1 substrates, and a few highly abundant 24 nt het-siRNAs were shown to be truncated and tailed in the *hen1* mutant (11, 13, 15). However, genomewide study of non-templated nucleotides in het-siRNAs is made more difficult by the heterogeneity inherent to het-siRNA biogenesis. Unlike miRNAs, where typically one or two major mature miRNA sequences accumulate, het-siRNAs from a given locus are much more sequence-diverse. This high diversity and general lack of a single dominant 'major' RNA species largely prevents approaches, such as miTRATA (18), that rely on identifying non-templated variants of a single abundant product. In this study, we take an alternative approach based on mismatches between aligned small RNAs to the corresponding reference genome to comprehensively profile single-nucleotide non-templated nucleotides in plant miRNAs and het-siRNAs.

## Materials and Methods

### Small RNA sequencing library preparation

*Arabidopsis thaliana* (ecotype Col-0) plants were grown at 21°C, with 16 h day/8h night. Inbred B73 maize plants were grown in a greenhouse at ~28°C. Total RNA from immature *Arabidopsis* inflorescence and fully expanded maize leaves was extracted using the miRNeasy Mini kit (Qiagen) per the manufacturer's instructions. Small RNA libraries were prepared using the TruSeq Small RNA kit (Illumina) per the manufacturer's instructions. Small RNA libraries were sequenced on HiSeq2500 (Illumina) with 50 nt read single-end runs. Raw data have been deposited at NCBI GEO under accession GSE79119 *(Arabidopsis* libraries) and GSE77657 (maize libraries).

### Source and processing of small RNA sequencing datasets

Sources and accessions of small RNA sequencing datasets analyzed in this study are in Table S1. For *A. thaliana* AGO4 immunoprecipitation datasets (raw accession numbers SRR189808, SRR189809, SRR189810, and SRR189811), 3' adapters were removed by Cutadapt (19) with options ‐a TCGTATGC ‐e 0.1 ‐0.5 ‐m 15. Trimmed reads were then aligned to the *A. thaliana* reference genome by ShortStack 3.3 (2, 3) with default settings. Other datasets were processed directly by ShortStack 3.3 with default settings. 3'-adapter sequences of all libraries and reference genomes used for alignment are shown in Table S1. The alignment settings retained an alignment containing a single mismatch only if there were no possible perfectly aligned positions for the read in question.

### Preparation of Simulated sRNA-seq Libraries

Simulation was accomplished through the use of the script sRNA-simulator.py (Supplemental script) with default settings. Selection of loci for simulated libraries used a hybrid approach based on prior annotations as well as real alignment profiles of sRNA-seq libraries, producing the three commonly-studied classes: miRNAs, het-siRNAs and trans-acting siRNAs (tasiRNAs). 15 *Arabidopsis*, 12 rice and 21 maize libraries (Table S2) were sourced for simulation, using the TAIR10, MSU 7 and AGPv3 reference genome assembly, respectively. Selected libraries were aligned to corresponding reference genome using Bowtie (20) with settings to map all locations for every read. High-confidence miRNA loci as listed in miRBase 21 (21) were used for miRNA simulation. Loci that were dominated by 23-24 nt or 21 nt RNAs were not used for miRNA simulation, and were considered candidates for het-siRNA and tasiRNA loci, respectively. Approximate 5 million reads were simulated from selected small RNA producing loci in each library. Thirty percent of simulated reads mimicked RNAs from miRNA loci, with the abundant 21 nt RNAs in a stranded miRNA/miRNA* pattern. Five percent of simulated reads mimicked tasiRNAs, with 21 nt RNAs arising from 125 nt long loci in a phased pattern on both strands. Het-siRNAs represented 65% of total reads, as 24 nt siRNAs were produced from 200-1000 nt long loci, which closely emulating het-siRNA processing *in vivo* (22, 23). Single nucleotide errors were randomly incorporated at a rate of one nucleotide per 10,000 reads. All loci produced a distribution of differently sized reads, mimicking DCL mis-processing. Once produced, simulated data were analyzed identically to real data.

### Analysis of non-templated nucleotides

BAM-formatted alignment files from ShortStack 3.3 (2, 3) output were used to analyze mismatched nucleotides. Non-mappers and secondary alignments were removed from the BAM files by SAMtools 1.1 (24) with options view ‐b ‐F 4 ‐F 256. The remaining reads were then intersected into high-confidence miRNA clusters (21) as listed in miRBase 21, ShortStack *de novo* annotated miRNA clusters, clusters dominated by 21 nt siRNAs, clusters dominated by 24 nt siRNAs, and clusters where less than 80% of the aligned reads were between 20 and 24 nts by Bedtools 2.19.1 (25). Most *de novo* annotated *MIRNA* loci by ShortStack overlapped prior miRBase annotations. *MIRNA* loci that were truly novel and were reproducible (called *de novo* by ShortStack from all analyzed sRNA-seq libraries in specific species) were shown in Table S3 and Supplemental Dataset 1. If a reference genome build used in this study was different from the one in miRBase 21, coordinates of high-confidence miRNA loci were updated by aligning the hairpin sequence from miRBase 21 to the appropriate genome build using BLAST 2.2.30 (26). Reads in each cluster were further grouped based on their length. The 'MD:Z' field from the alignment lines was used to determine positions of bases with mismatches between the read and the reference genome. Fractions of reads with mismatches in each cluster class were calculated. To analyze mismatch frequencies in 24 nt-dominated clusters, biological replicates were merged as a single library. Reads with or without mismatched nucleotides within the same size group were processed by WebLogo 3.4 (27) with options ‐‐format pdf ‐‐yaxis 1 ‐‐color-scheme classic.

## Results

### Elevated mismatch rates at the 3' ends of genome-aligned plant miRNAs and siRNAs are due to non-templated nucleotides

To study global patterns of non-templated nucleotides in plant small RNAs, we first analyzed small RNA deep sequencing (sRNA-seq) libraries prepared from wild-type *Arabidopsis*, maize, and rice tissues (28). After adapter trimming, reads with lengths of 15 nt or longer were aligned to their corresponding genomes allowing no more than one mismatch. Alignments with single mismatches were allowed only if there were no perfectly-matched alignments for a given read. Each alignment with a single mismatch could potentially arise from a non-templated nucleotide, sequencing error, software error, a single-nucleotide polymorphism (SNP) between the specimen sampled for sRNA-seq and the reference genome assembly, or an error in the reference genome assembly itself. We observed elevated rates of mismatches at the 3'-most nucleotide regardless of RNA sizes and plant species (Figure 1A). While mismatches at 3' positions were always elevated, the rates also varied by RNA size and species (Figure 1A). The position-specific nature of this pattern rules out SNPs and reference genome assembly errors as major contributors because there is no obvious reason that these situations would consistently result in mismatches predominantly at the 3'-most nucleotides of the reads.

**Figure 1.**
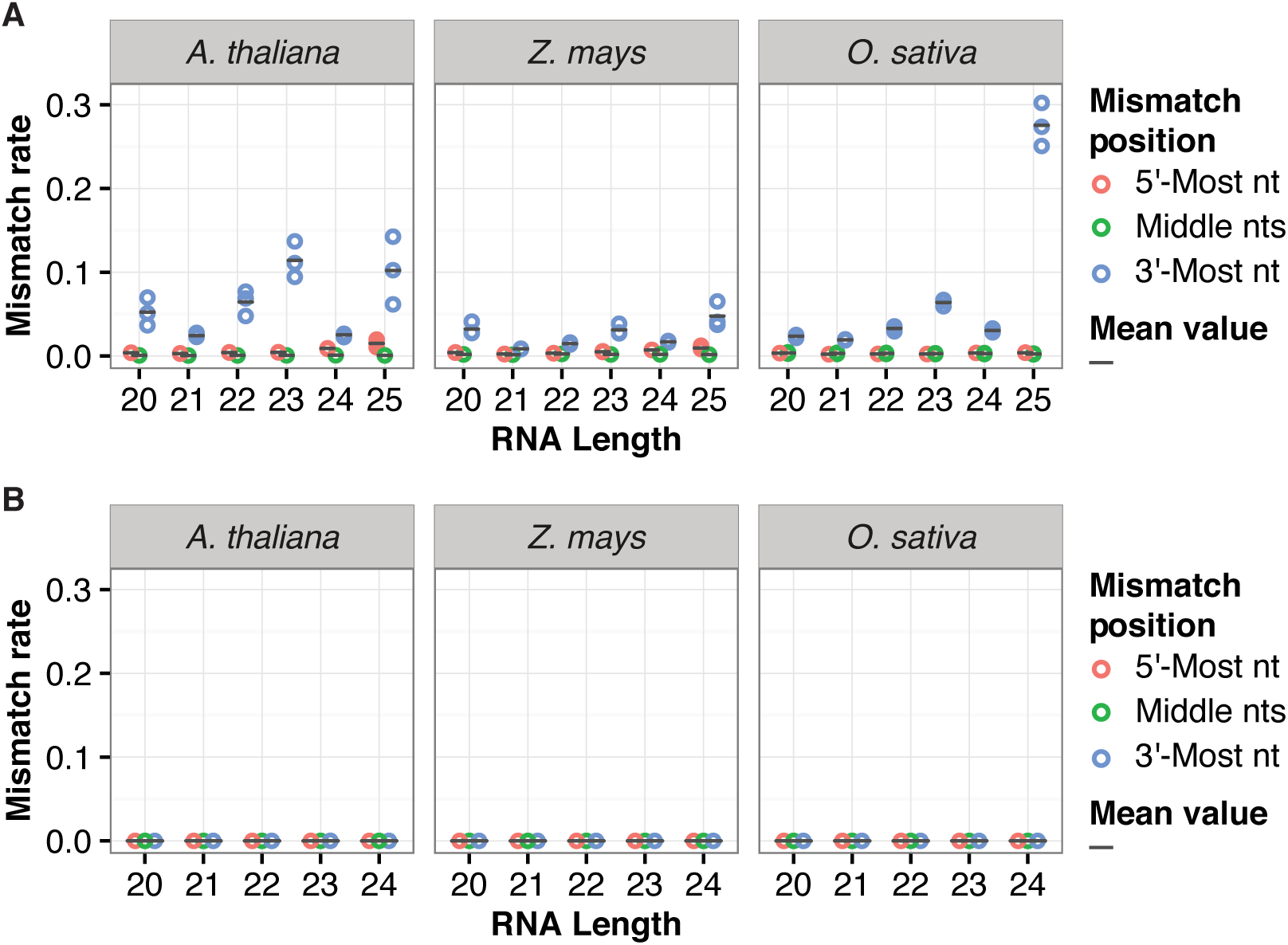
Elevated mismatch rates at the 3'-most positions of genome-aligned plant small RNAs. A. Mismatch rates for reference-genome aligned sRNA-seq reads of the indicated sizes and species. Rates for the 5'-most nt, all middle nts, and the 3'-most nt are plotted separately. Multiple data points for the same species and RNA size indicate values from different replicate sRNA-seq libraries. Circles: biological replicates. Bars: mean value. Libraries that were analyzed are listed in Table S1. B. Mismatch rates for reference-genome aligned sRNA-seq reads from simulated datasets.

To test whether the elevated 3' mismatch rates were caused by software errors, we first examined simulated sRNA-seq datasets. 15 simulated *Arabidopsis* small RNA libraries, 21 simulated maize small RNA libraries, and 12 simulated rice small RNA libraries were aligned to their corresponding reference genomes. The simulated data included singlenucleotide errors at random positions at a rate of 0.01%. We didn't observe elevated mismatch rates on the 3' ends of the simulated libraries (Figure 1B). We then tested whether the elevated 3' mismatch rates were due to software issues associated with 3' adapter trimming; the simulated data did not include 3' adapters and as such weren't trimmed. We trimmed 3' adapter sequences of the same real *Arabidopsis* sRNA-seq datasets by using Cutadapt 1.8.3 (19) instead of ShortStack 3.3 (2, 3). Elevated rates of 3' mismatches were still observed for all size groups of small RNAs (Figure S1). These results indicate that the high frequencies of 3' mismatches seen in plant sRNA-seq data are not likely due to a systemic software artifact.

We next tested whether the pattern of 3'-mismatches was due to general sequencing errors. To do this, we grouped aligned *Arabidopsis* sRNA-seq reads into genomic clusters and classified the clusters. Two non-mutually exclusive groups of miRNAs were identified: Those from our *de novo* analysis, and those listed as high-confidence loci in miRBase 21. Three groups of non-miRNA loci were also analyzed: Loci dominated by 21 nt siRNAs, loci dominated by 24 nt siRNAs (which we presume are mostly het-siRNAs), and loci where less than 80% of the aligned reads were 20-24 nts in length (Which we termed N loci). The N loci likely represent degraded tRNAs, rRNAs, mRNAs, and other cellular RNAs that are not related to the DCL / AGO regulatory system. The reads aligned to the N loci did not have high rates of 3'-mismatches (Figure 2A). High rates of 3'-mismatches were confined to certain RNA sizes within miRNA, 21 nt loci, and 24 nt loci (Figure 2A). These trends were not unique to our sRNA-seq libraries; the same analysis procedure applied to previously published *Arabidopsis* sRNA-seq data (Table S1) gave similar results (Figure S2). The specificity of the high 3'-mismatch rate for miRNAs and siRNA loci, but not for degraded RNA loci, argues against sequencing errors as a major contributor. Based on these analyses, we conclude that the high 3'-mismatch rates seen for miRNAs and siRNAs in genome-aligned plant sRNA-seq data are due to the presence of non-templated nucleotides *in vivo*.

**Figure 2.**
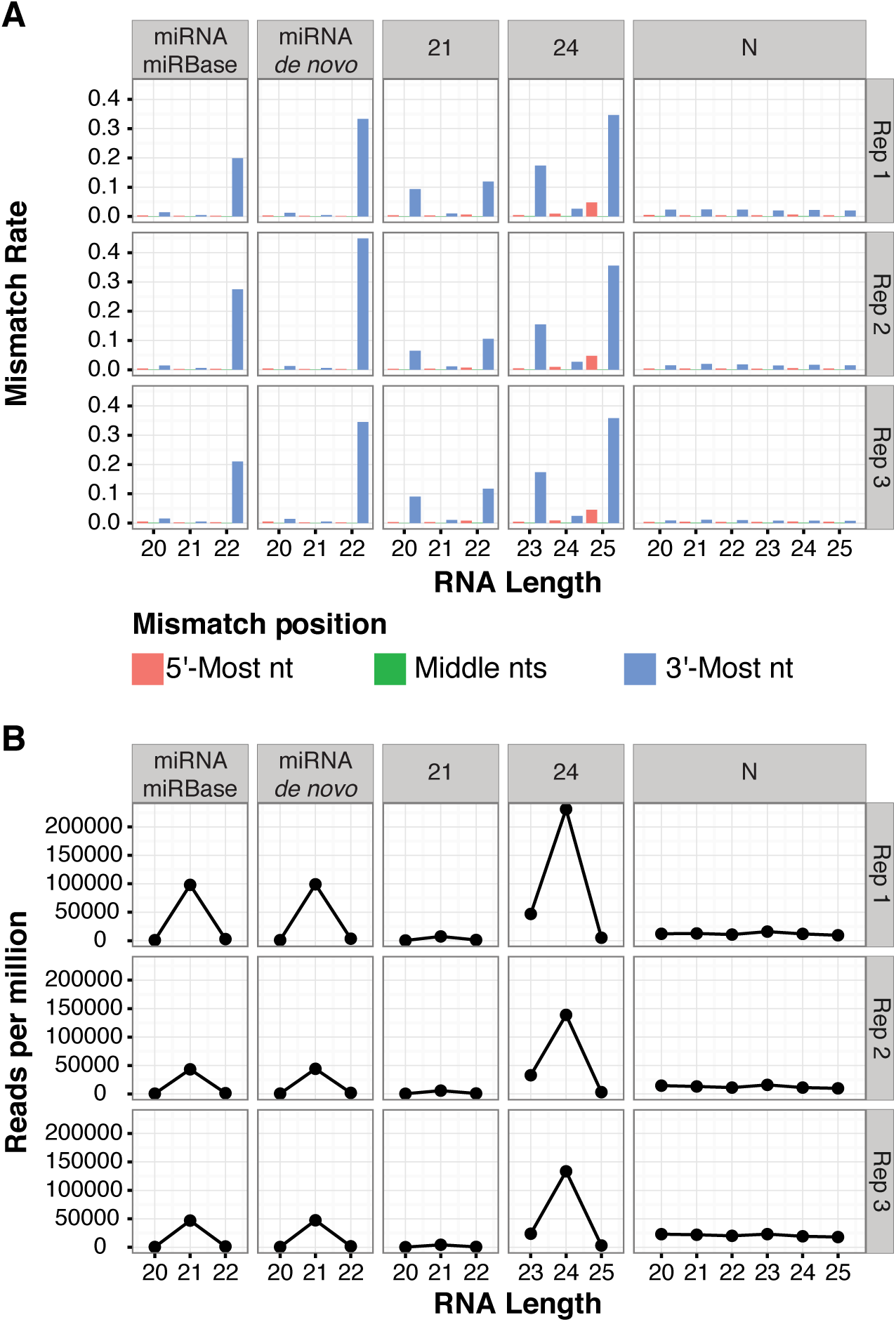
'Off-sized' miRNAs and siRNAs have high rates of 3' mismatches in *Arabidopsis*. A. Mismatch rates for reference-aligned *Arabidopsis* sRNA-seq reads of the indicated clusters and sizes. 21: siRNA loci dominated by 21 nt RNAs. 24: siRNA loci dominated by 24 nt siRNAs (presumed het-siRNA loci). N: loci where less than 80% of reads were between 20-24 nts, and are thus presumed to not be miRNA or siRNA loci. miRNAmiRBase: High-confidence *MIRNA* loci from miRBase version 21. miRNA-de novo: *MIRNA* loci annotated by automated software. Note that most *de novo* annotated *MIRNA* loci overlapped prior miRBase annotations. *Arabidopsis* libraries that were analyzed are listed in Table S1. B. As in A, except showing normalized accumulation levels of small RNAs.

### 'Off-sized' het-siRNAs and miRNAs often have higher rates of 3' end non-templated nucleotides

In *Arabidopsis*, most abundant miRNAs are 21 nts in length (Figure 2B). Relatively low rates of non-templated 3' nucleotides were observed for the predominant sizes of miRNAs, 21 nt siRNAs, and 24 nt siRNAs (Figure 2A). By contrast, 'off-sized' small RNAs showed higher rates of 3' non-templated nucleotides, with the small RNAs one nucleotide longer than the predominant size usually having the highest rates (Figure 2A). For example, about 35.3% of 25 nt siRNAs and about 16.7% of 23 nt siRNAs aligned to 24 nt-dominated clusters had a 3' end non-templated nucleotide. In contrast, only about 2.6% of 24 nt siRNAs aligned to 24 nt-dominated clusters had a 3' end non-templated nucleotide. These trends were also apparent in other previously published *Arabidopsis* sRNA-seq data (Figure S2).

We further studied whether the elevated rate of 3' end non-templated nucleotides for 'off-sized' small RNA could be observed in other plant species. 17 sRNA-seq libraries from wild-type specimens of *Solanum lycopersicum* (29, 30), *Medicago truncatula* (31), *Zea mays* (this study), *Oryza sativa* (28) and *Physcomitrella patens* (32), along with 9 sRNA-seq libraries from *Arabidopsis thaliana* (same libraries as in Figure 2 and S2), were analyzed (Table S1). 22 nt siRNAs from 24 nt-dominated siRNA clusters in *Physcomitrella* were included in the analysis, because 23 nt siRNAs and 24 nt siRNAs from 24 nt-dominated siRNA clusters in *Physcomitrella* are similarly abundant (32). Similar to *Arabidopsis*, we observed that small RNAs of the predominant size for their locus type usually have low rates of 3' non-templated nucleotides in all species examined (Figure 3). miRNAs and siRNAs with one nucleotide longer than predominant sizes in all clusters had higher rates of 3' end non-templated nucleotides, with 25 nt siRNAs from 24 nt-dominated clusters usually having the highest levels (Figure 3). As in *Arabidopsis*, RNAs aligned to N clusters (which are unlikely to be miRNAs or siRNAs) usually did not have elevated rates of 3' non-templated nucleotides (Figure 3). We noted that 25 nt RNAs from *Oryza sativa* N clusters also had elevated rates of 3'-mismatches (Figure 3). We speculate that this could be due to mis-classification of some clusters that are truly 24 nt siRNA dominated clusters as N clusters in this species.

**Figure 3.**
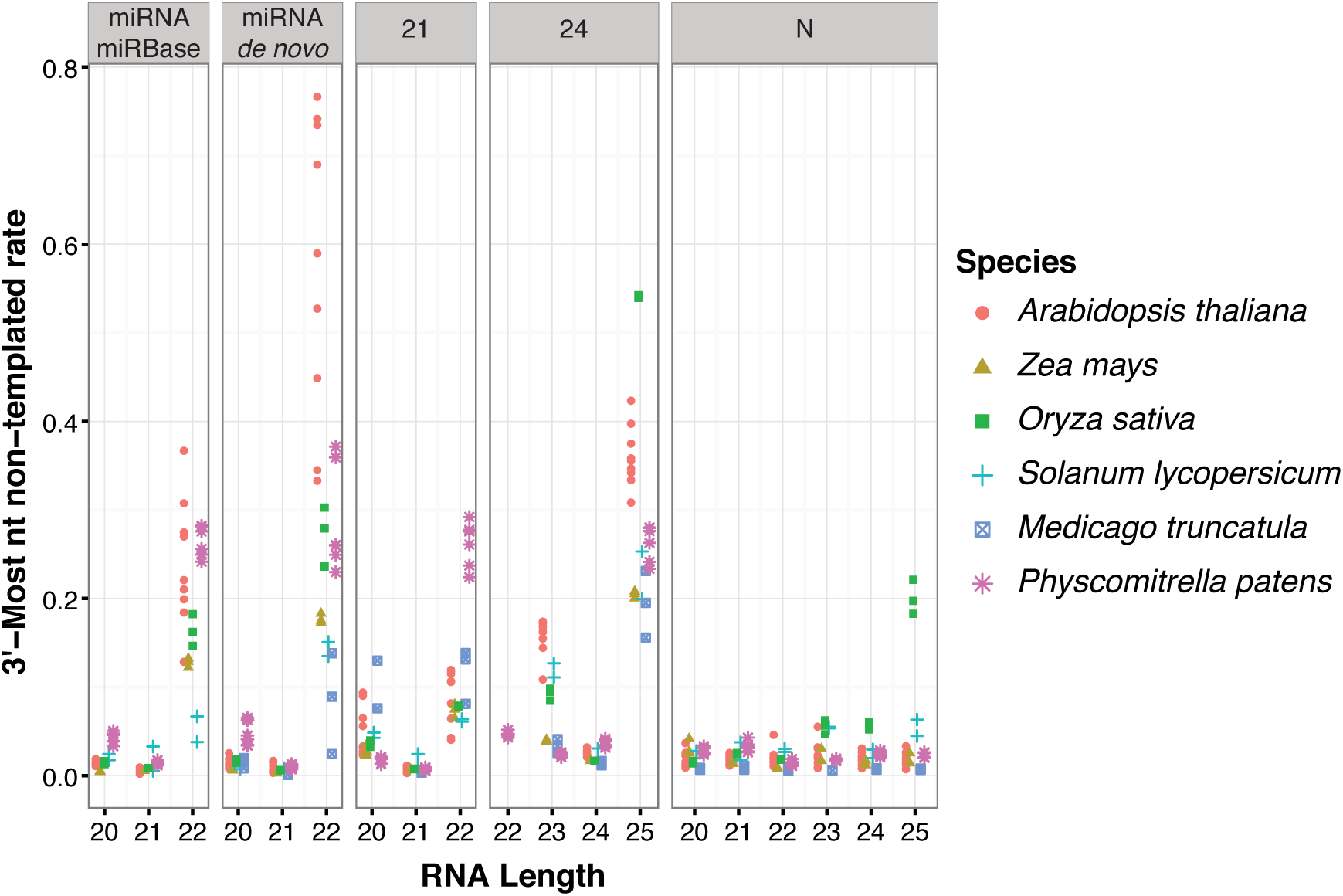
'Off-sized' miRNAs and siRNAs have high rates of 3' end non-templated nucleotides in several plant species. Mismatch rates for reference-aligned sRNA-seq reads of the indicated species, clusters, and sizes. 21: siRNA loci dominated by 21 nt RNAs. 24: siRNA loci dominated by 24 nt siRNAs (presumed het-siRNA loci). N: loci where less than 80% of reads were between 20-24 nts, and are thus presumed to not be miRNA or siRNA loci. miRNA-miRBase: High-confidence *MIRNA* loci from miRBase version 21. miRNA-de *novo: MIRNA* loci annotated by automated software. Note that most *de novo* annotated *MIRNA* loci overlapped prior miRBase annotations. Libraries that were analyzed are listed in Table S1.

### *Arabidopsis* 23 nt siRNAs from 24 nt-dominated clusters have a unique pattern of 3' non-templated nucleotides

The 3'-most nucleotides of plant siRNAs and miRNAs are 2'-*O*-methylated by the HEN1 methyltransferase (10, 11). In the absence of HEN1 activity, miRNAs and siRNAs are subject to 3'-uridylation (11). Thus, one hypothesis that could explain our observations is that HEN1 disfavors the 'off-sized' miRNAs and siRNAs, rendering them unmethylated and thus susceptible to 3'-end uridylation in the wild-type. Consistent with this hypothesis, U was a very frequent 3' non-templated nucleotide of *Arabidopsis* 25 nt siRNAs from 24 nt-dominated clusters (Figure 4). These 25 nt siRNAs also tended to have a 5'-A, as do canonical 24 nt het-siRNAs (Figure 4). However, 23 nt siRNAs from 24 nt-dominated clusters had a very different pattern. The non-templated 3'-most nucleotide for 23 nt siRNAs tended to be an A, while there is no strong tendency to have the canonical 5'-A (Figure 4). Surprisingly, position 22 of the 23 nt non-templated siRNAs has a very strong tendency to be a U or C (Figure 4). This pattern was also present in maize, rice, *Medicago truncatula*, and tomato (Figure S3). The U or C at position 22 is templated by the genome, suggesting that the 23 nt siRNAs with a 3'-mismatch occur when templated at specific genomic locations. The distinct preference of 3' non-templated nucleotides indicates that distinct mechanisms are likely to underlie their deposition on 23 nt and 25 nt siRNAs from *Arabidopsis* 24 nt-dominated loci.

**Figure 4.**
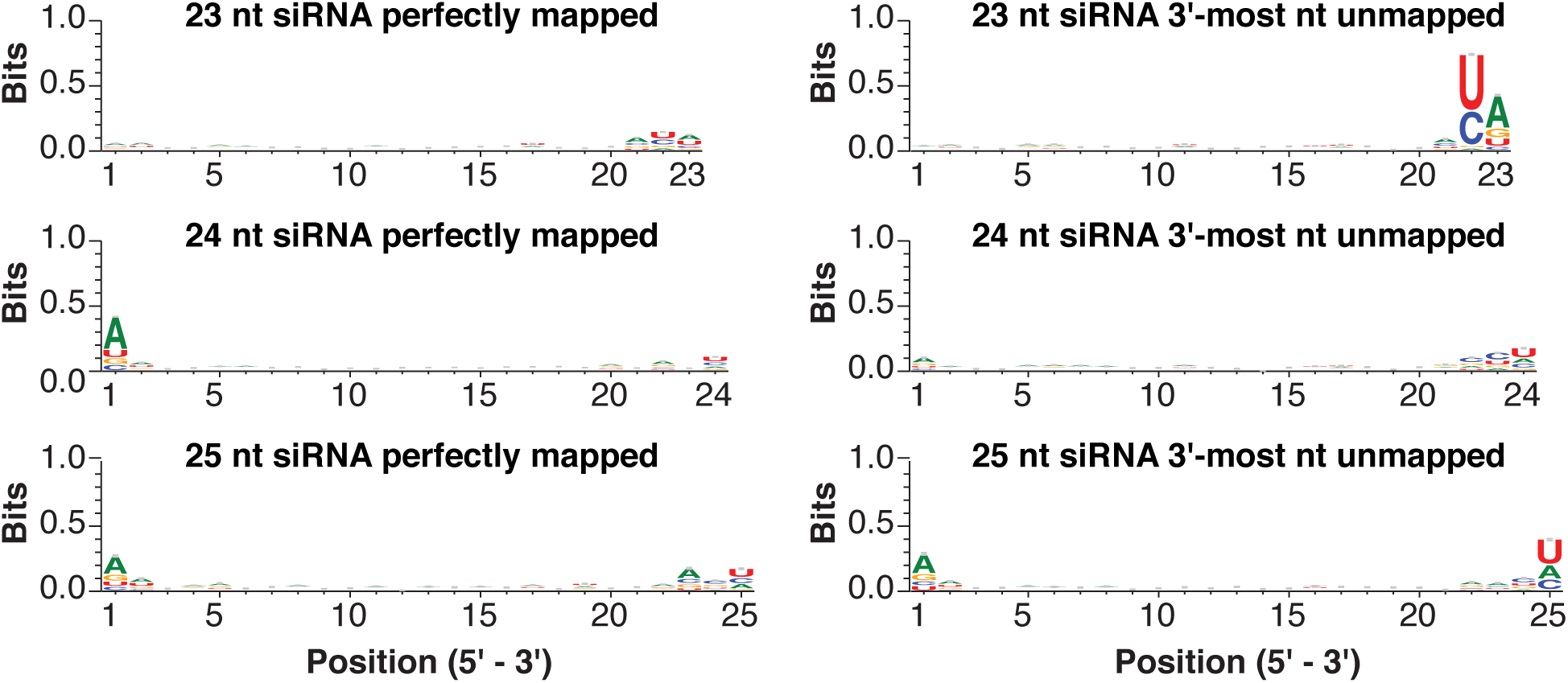
Sequence features of siRNAs arising from *Arabidopsis* 24 nt-dominated siRNA loci. Sequence logos for the indicated siRNAs aligned to 24 nt-dominated siRNA loci. Three biological replicate sRNA-seq libraries from *Arabidopsis* inflorescence (Table S1) were merged for this analysis. Sequence features were analyzed by WebLogo 3.4.

We next examined *Arabidopsis* sRNA-seq data from *hen1* mutants, as well as the 3' uridylase mutants *hesol* and *urtl* (15). As expected, increased levels of 3' non-templated uridylation were observed in the *hen1* mutant for all sizes of siRNAs arising from 24 nt-dominated clusters (Figure 5A). This uridylation was suppressed in *hen1/heso1* and especially *hen1/hesol/urtl* backgrounds. Similar trends of *HESOl-*and *URTl-dependent* 3' uridylation in the *hen1* background were observed for miRNAs (Figure 5B). However, the 3' non-templated adenylation of 23 nt siRNAs from 24 nt-dominated clusters decreased, not increased, in the *hen1* mutant (Figure 5A). Together with the analysis of sequence motifs, these data suggest that the frequent 23 nt siRNAs with 3' non-templated A residues are produced by a mechanism distinct from the currently understood HESO1‐ and URT1-dependent terminal transferases.

**Figure 5.**
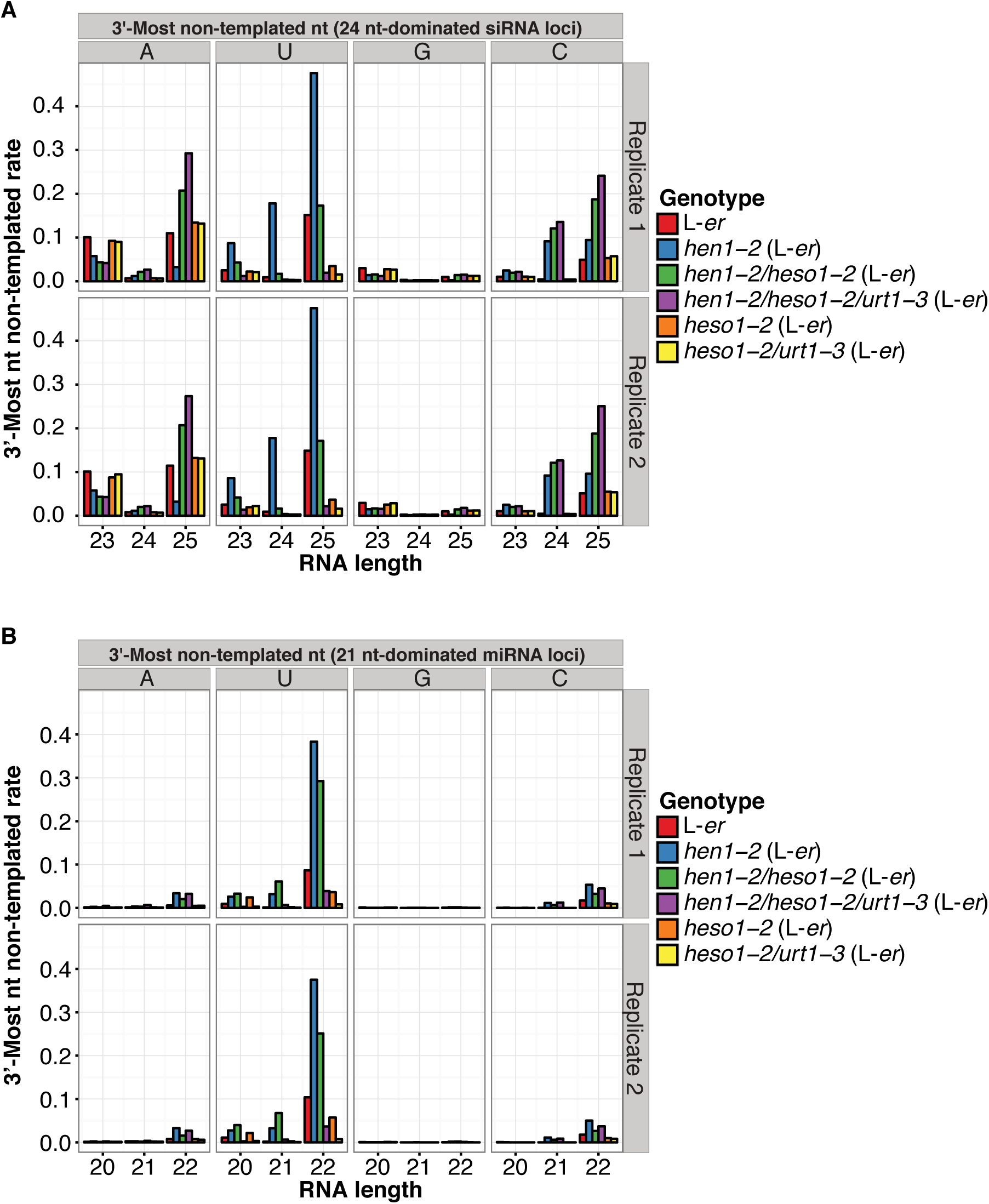
3'-most non-templated adenines in 23 nt siRNAs arising from 24 ntdominated siRNA clusters are not dependent on HESO1 and URT1. A. Rates of *Arabidopsis* 3' non-templated nucleotides in siRNAs aligned to 24 ntdominated loci by siRNA size, 3'-most non-templated nucleotide, and genetic background. Data from Wang *et al*. (15) (Table S1). *L-er*. Landsberg *erecta* ecotype. B. As in A except for miRBase high-confidence miRNA clusters.

24 nt siRNAs in *Arabidopsis* are mostly generated by the Dicer-Like enzyme DCL3 (33, 34). However, DCL2 and DCL4 can also contribute 22 nt and 21 nt siRNAs, respectively, at loci dominated by 24 nt siRNAs (34). Therefore, we tested whether the 23 nt siRNAs with a 3'-most non-templated nucleotide were derived from tailing of DCL2-dependent 22 nt siRNAs by examining sRNA-seq data from a *dcl2/dcl4* double null mutant. 23 nt siRNAs with a 3'-most non-templated nucleotide were still present at 24 nt-dominated siRNA loci in the *dcl2/dcl4* background, and had the same pattern of nucleotide biases at their 3' ends (Figure S4). Thus we conclude that the population of 23 nt siRNAs with a 3' non-templated nucleotide is not generally produced by DCL2 or DCL4.

### 23 nt siRNAs with a 3' non-templated nucleotide are infrequently bound to AGO4

We then studied whether siRNAs with a 3' non-templated nucleotide from 24 nt-dominated clusters were loaded into AGO4 by analyzing *Arabidopsis* AGO4 immunoprecipitation sRNA libraries (35). The rate of 3' non-templated nucleotides in AGO4-immunoprecipitated 24 nt siRNAs was low (2.4%, Figure 6A). In contrast, 24.8% of AGO4-immunoprecipitated 25 nt siRNAs were carrying a 3' non-templated nucleotide (Figure 6A). Interestingly, compared to total RNA, a much reduced percentage of AGO4associated 23 nt siRNAs had 3' non-templated nucleotides (16.7% in total RNA vs. 3.9% in the AGO4 immunoprecipitates; Figures 2A, 6A). All AGO4-associated siRNAs had a strong 5' adenine preference regardless of size and the presence of a 3' end non-templated nucleotide (Figures 6C-D). The strong 5'-A preference was not observed for 23 nt siRNAs with a 3' non-templated nucleotide in total RNA (Figure 4). Thus, we speculate that the known selectivity of AGO4 for 5'-A containing siRNAs (36) reduces the loading of 23 nt siRNAs with 3' non-templated nucleotides onto AGO4. URT1 and HESO1 can act upon AGO-bound small RNAs (14, 16). The fact that 25 nt siRNAs with a nontemplated 3' nucleotide are bound to AGO4 at robust frequencies, coupled with their tendency to have 5'-As and a U as the non-templated 3' nucleotide, suggests they are HESO1 and/or URT1 substrates. However, the 23 nt siRNAs with a non-templated 3' nucleotide are not frequently bound to AGO4, suggesting that their 3' non-templated nucleotides are likely to be added at a step prior to AGO4 loading. However, it is also possible that the 23 nt siRNAs with a non-templated 3' nucleotide might instead be bound to an Argonaute other than AGO4.

**Figure 6.**
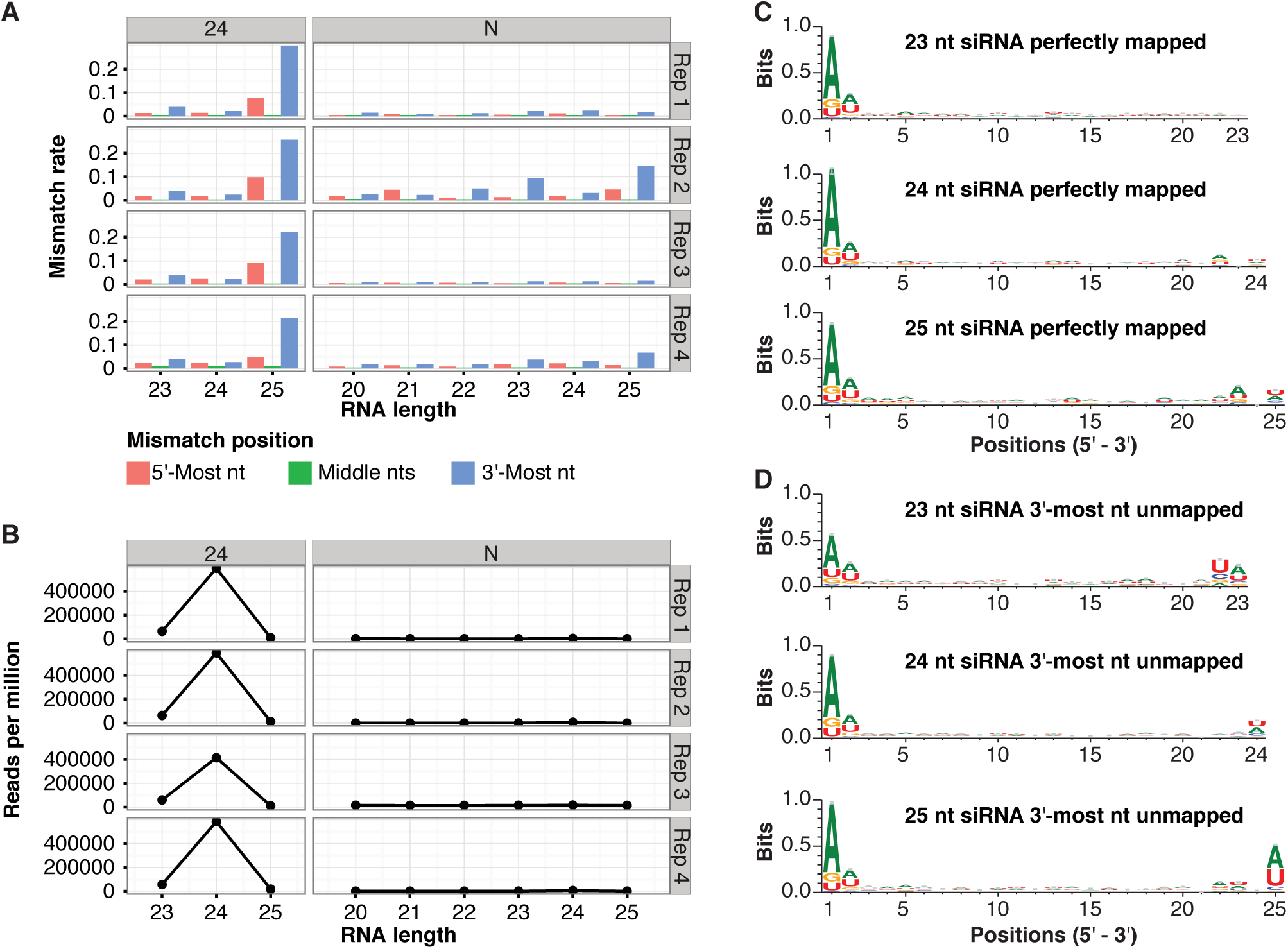
Rates of non-templated nucleotides in siRNAs co-immunoprecipitated with *Arabidopsis* AGO4. A. Rates of non-templated nucleotides in AGO4-immunoprecipitated RNA, indicated by cluster type and RNA sizes. 24: 24 nt-dominated siRNA loci. N: loci where less than 80% of the aligned reads were 20-24 nts in length. Data from Wang *et al*. (35) (Table S1). B. As in A, except showing normalized accumulation levels of small RNAs. C. Sequence logos for perfectly aligned siRNAs arising from 24 nt-dominated siRNA clusters. The four *Arabidopsis* AGO4 immunoprecipitation sRNA-seq libraries from Wang *et al*. (35) (Table S1) were merged for this analysis. Sequence features were analyzed by WebLogo 3.4. D. As in C, except for reads with a 3'-most non-templated nucleotide.

### 23 nt siRNAs from 24 nt-dominated clusters are mostly not 3'‐ or 5'-truncated variants of 24 nt siRNAs

We next examined whether the 23 nt siRNAs from 24 nt-dominated clusters are frequently 3' or 5' truncated variants of the more prevalent 24 nt siRNAs. The frequencies at which the 5' or 3' ends of *Arabidopsis* 23 nt siRNAs overlap with 24 nt siRNAs from the same clusters were calculated. Lowly expressed clusters (those with less than 20 aligned 24 nt siRNAs) were excluded from analysis. The 23 nt siRNAs were infrequently 5'-truncated variants (median values ~25%) and 3'-truncated variants (median value ~12%) of 24 nt siRNAs (Figure 7A). 23 nt siRNAs with 3'-most non-templated nucleotide are even less frequently 5'‐ or 3'-truncated variants of 24 nt siRNAs (median values of 0%) (Figure 7A). 25 nt siRNAs, however, showed a higher tendency to be 3'‐ tailing variants (median values ~35%, for RNAs with 3'-most non-templated nucleotide), but not 5'-tailing variants (median values of 0%) of 24 nt siRNAs. As a control, we examined 20 nt and 22 nt RNAs arising from high confidence miRNA loci in *Arabidopsis*. The majority of these 20 nt and 22 nt RNAs were 3'-truncation or 3'-tailing variants of a more abundant 21 nt RNA (most often the major mature miRNA) (Figure 7B). These observations suggest that the 23 nt siRNAs present in *Arabidopsis* 24 nt-dominated loci are generally not 3'‐ or 5'-truncations of 24 nt siRNAs, whereas 25 nt siRNAs are more likely to be 3'-tailings of 24 nt siRNAs.

**Figure 7.**
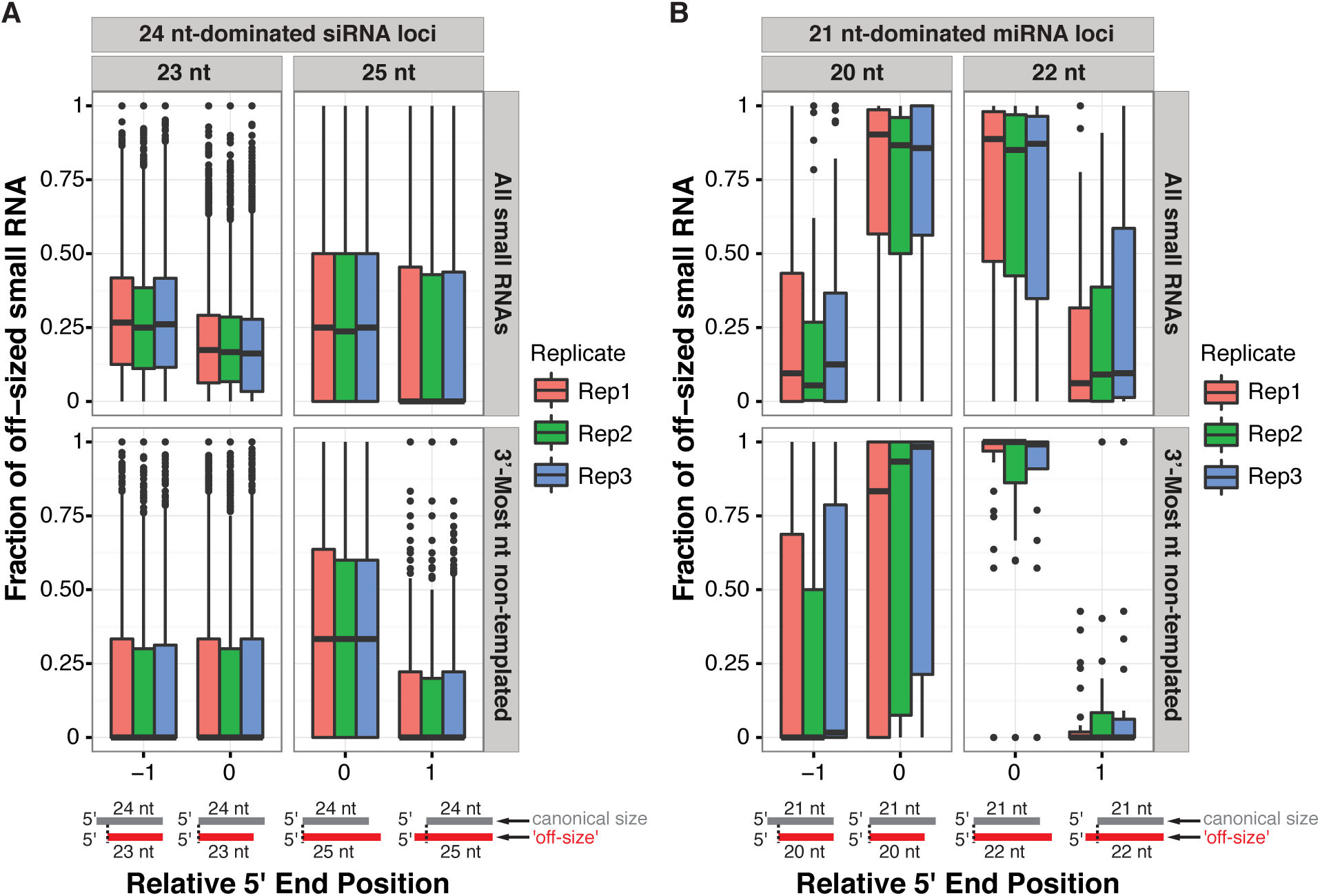
Analysis of 5'‐ and 3'-truncations and tailings for 'off-sized' *Arabidopsis* small RNAs. A. Fractions of 23 and 25 nt siRNAs which shared coincident 5' or 3' ends with 24 nt siRNAs in robustly expressed (>= 20 24nt reads) 24 nt-dominated siRNA clusters. Boxplots show medians (horizontal lines), the 1st-3rd quartile range (boxes), other data out to 1.5 times the interquartile range (whiskers) and outliers (dots). Data from *Arabidopsis* inflorescences (Table S1). Illustrations below depict different possible relative 5'-end relationships between the canonical sized RNAs (gray) and off-sized RNAs (red). B. As in A, except for 20 nt and 22 nt RNAs aligned to high-confidence miRNA loci compared to 21 nt miRNAs.

### Further properties of 23 nt siRNAs with a 3'-most non-templated nucleotide

In the absence of DCL proteins, RNAs longer than 24 nts accumulate from *Arabidopsis* het-siRNA loci (22, 23, 37-39). These RNAs, termed P4R2 RNAs because of their dependence on both DNA-dependent RNA polymerase IV (Pol IV) and RNA-dependent RNA polymerase 2 (RDR2), are likely to be the direct precursors for DCL3-catalyzed production of het-siRNAs. Many P4R2 RNAs have 3' non-templated nucleotides which are thought to predominantly be located on the Pol IV-transcribed strand (22, 23, 38). Zhai et al (23) reported that cytosines were enriched in the template DNA sequence at the 3'-most non-templated positions of P4R2 RNAs and hypothesized that 5-methyl cytosines promote Pol IV transcriptional termination and addition of a non-templated nucleotide. We analyzed the DNA nucleotide frequencies at positions corresponding to the 3' non-templated nt in siRNAs from 24 nt-dominated loci. Cytosines were strongly enriched at the 3'-most non-templated positions of 23 nt siRNAs, but not in other size groups (Figure 8A). We next tested if 23 nt siRNAs with a 3'-most non-templated nucleotide were frequently the passenger strand siRNA for 24 nt siRNAs. This analysis assumed a 2 nt offset of the 5' end (consistent with DCL catalysis), and a 1 nt offeset on the 3' end (assuming addition of a non-templated nt to the end of an originally blunt-ended dsRNA). These arrangements were quite rare (Figure 8B). If the hypothesis that canonical 24 nt siRNAs mostly derive from the 5'-ends of the Pol IV transcribed strands of P4R2 RNAs is correct (23), then this observation suggests that the 23 nt siRNAs with a 3' non-templated nucleotide do not frequently originate from the RDR2-transcribed strand. Overall, these observations are consistent with the hypothesis that these 23 nt siRNAs are more frequently derived from the 3' ends of the Pol IV-transcribed strand of P4R2 RNAs as opposed to the 3' ends of the RDR2-transcribed strands.

**Figure 8.**
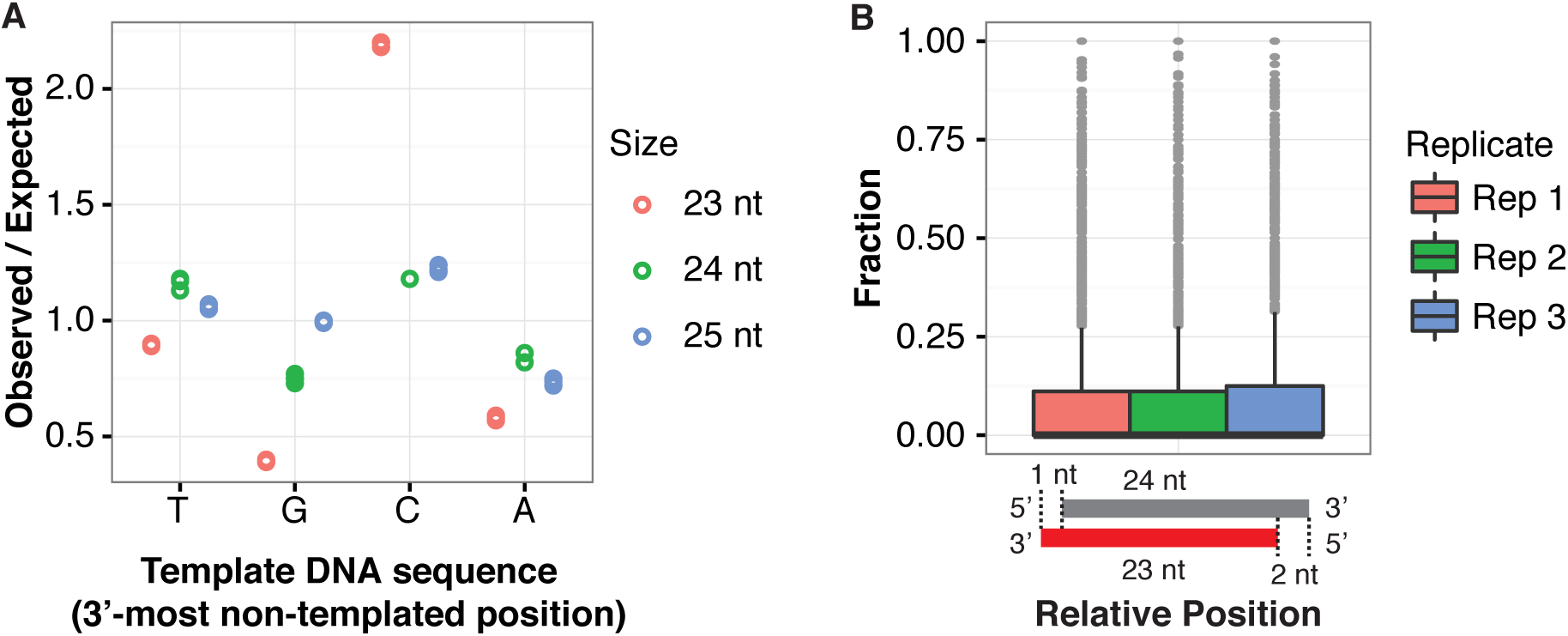
Further properties of 23 nt siRNAs with a 3'-most non-templated nucleotide. A. Enrichment/depletion analysis of genomic nucleotides corresponding to the 3'-most non-templated positions of 23 nt, 24 nt and 25 nt siRNAs arising from 24 nt-dominated siRNA clusters. Three wild-type *Arabidopsis* sRNA-seq libraries (Table S1) were analyzed. Biological replicates are represented by circles. B. Frequency of 23 nt siRNAs with a 3’-most non-templated nucleotide that correspond to a reverse-complemented 24 nt siRNA in 24 nt-dominated siRNA clusters. Three wildtype *Arabidopsis* sRNA-seq libraries were analyzed (Table S1). Lowly expressed clusters (those with less than 20 aligned 24 nt siRNAs) were excluded from analysis. Boxplots show medians (horizontal lines), the 1st-3rd quartile range (boxes), other data out to 1.5 times the interquartile range (whiskers) and outliers (dots).

## Discussion

Previous analyses of non-templated nucleotides in plant small RNAs have analyzed minor variants of highly abundant RNAs that do not contain any non-templated nucleotides (16, 18). This allows thorough analysis of both truncation and tailing variants. This approach is well-suited for miRNAs, for which a single dominant RNA sequence accumulates to high levels, allowing relatively easy identification of truncated and tailed variants. However, this method is ill-suited to characterize plant het-siRNAs because, unlike miRNAs, het-siRNA clusters are composed of multiple distinct siRNAs rather than a single dominant product. Here we show the utility of an alternative method to examine non-templated nucleotides in plant sRNA-seq data that can be applied to hetsiRNAs: genome alignment. By allowing valid alignments to contain one mismatch to the reference genome (provided that there are no zero-mismatch alignments for the read), we can capture RNAs with a single non-templated nucleotide. Importantly, control experiments rule out alternative explanations (software errors, sequencing errors, SNPs, and errors in the reference genome assembly) for the patterns of mismatched nucleotides.

Data from multiple plant species indicate that 'off-sized' 25 nt and 23 nt siRNAs that arise from clusters dominated by 24 nt siRNAs have high rates of single 3' non-templated nucleotides (Figure 3). We presume that most of these loci are het-siRNA loci, based on their dominant production of 24 nt siRNAs. Our data suggest that the 25 nt siRNAs with 3' non-templated nucleotides are added after AGO4 binding by the URT1 and/or HESO1 uridyltransferases (Figure 9). Supporting this hypothesis are the observations that the 25 nt siRNAs tend to have a 5'-A like the canonical 24 nt siRNAs (Figure 4), tend to have a U residue as the 3' non-templated residue (Figure 4), are increased in the *hen1* background in a *HESO1* and/or *URT1-dependent* manner (Figure 5), and are found associated with AGO4 (Figure 6). In contrast, the 'off-sized' 23 nt siRNAs with a 3' non-templated nucleotide had none of these features. Thus, we hypothesize that the 23 nt siRNAs with a 3' non-templated nucleotide are modified at a step prior to AGO4 binding in a manner insensitive to the presence or absence of the 2'-O-methyl modification deposited by HEN1 (Figure 9). It is worth noting that 25 nt siRNAs are rare at het-siRNA loci, but 23 nt siRNAs accumulate to significant levels in *Arabidopsis* (Figures 2B, S2).

**Figure 9.**
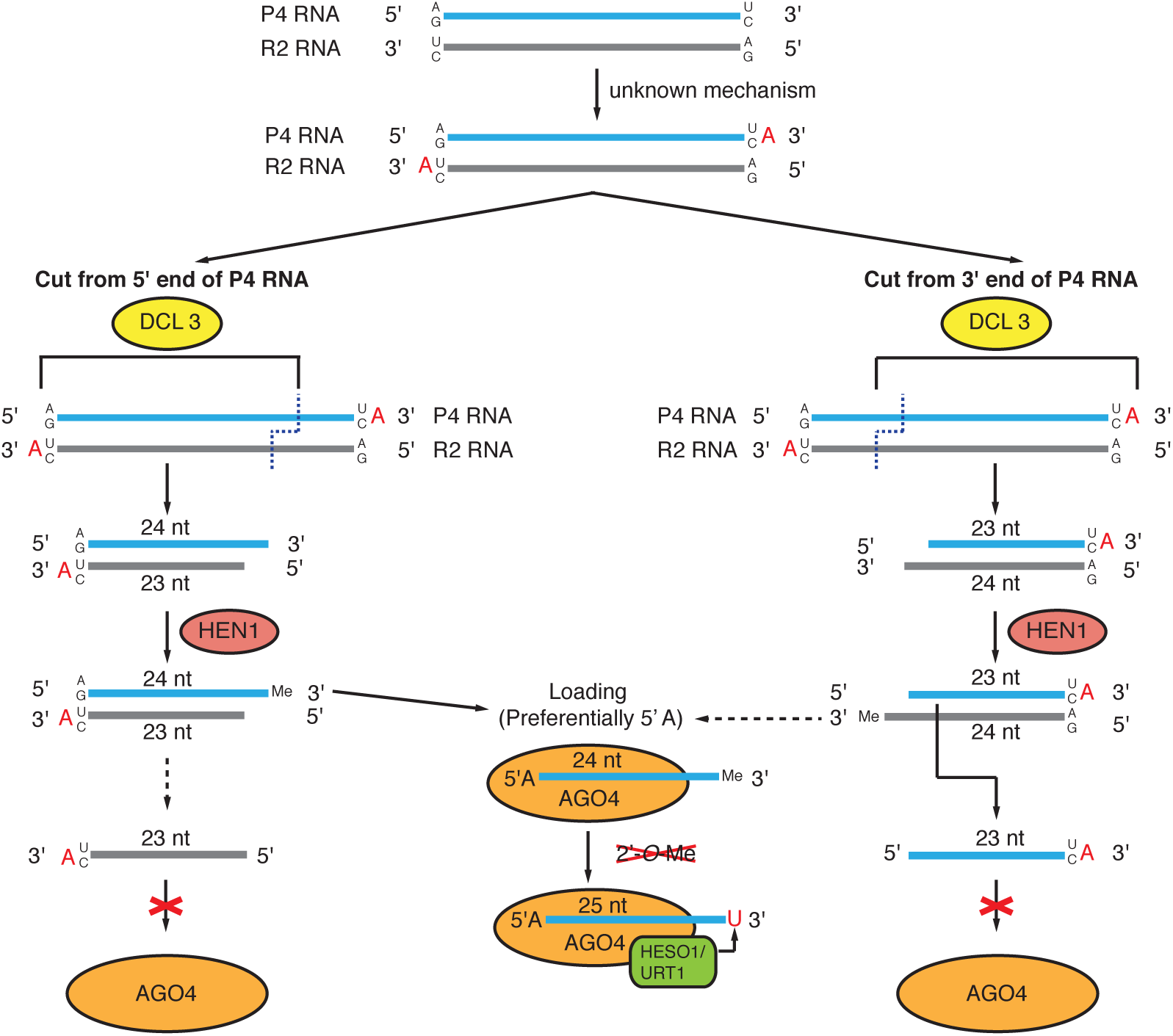
Model of 3' non-templated nucleotides in 'off-sized' siRNAs arising from 24 nt-dominated siRNA clusters. >Model of 3' non-templated nucleotides in 'off-sized' siRNAs arising from 24 ntdominated siRNA clusters. Non-templated nucleotides are indicated by red letters.

In *Physcomitrella patens*, 23 nt siRNAs accumulate to even higher levels at most het-siRNA loci (32, 40).

How might the 23 nt siRNAs with a 3' non-templated nucleotide arise? P4R2 RNAs tend to have a U or C at their 3' ends *in vivo* (22, 38), and *in vitro* transcription by Pol IV also results in RNAs with a strong enrichment of U or C at the 3' position (22). Pol IV transcribed RNAs also tend to begin with an A or G at the 5' most nucleotide (22, 23, 38), which means the 3' ends of the RDR2-transcribed second strands will also tend to be U or C. P4R2 RNAs also have a high frequency of 3' non-templated nucleotides (22, 23, 38). All of these data are consistent with a hypothesis where the 23 nt siRNAs with a 3' non-templated nucleotide result from DCL catalysis of a P4R2 precursor RNA that has a non-templated nucleotide (Figure 9). It is difficult at present to say whether the Pol IV-transcribed strand or the RDR2-transcribed strand contributes more to the pool of these 23 nt siRNAs. However, the observation that the corresponding DNA positions of the non-templated nucleotides tend to be C (Figure 8A), and that the 23 nt siRNA do not tend to be reverse complements of 24 nt siRNAs (Figure 8B) suggest they might more frequently derive from the Pol IV-transcribed strand. These inferences are based on the previous suggestions that the non-templated residues on Pol IV-transcribed RNAs tend to align with C's in the genome, and that the canonical 24 nt siRNAs are most often derived from the 5' ends of Pol IV-transcribed RNAs (23). Further testing of this hypothesis may shed light on the modes of DCL protein action and biogenesis of het-siRNAs in plants.

## Acknowledgements

We thank Craig Praul for small RNA-seq library sequencing, Qikun Liu, Quentin Gouil, Jake Harris, Jixian Zhai, Steven Jacobsen, and Xuemei Chen for helpful discussions and comments, and all members of the Axtell Lab for constructive comments on this work.

## References

1. Axtell,M.J. (2013) Classification and comparison of small RNAs from plants. Annu Rev Plant Biol, 64, 137–159.

2. Axtell,M.J. (2013) ShortStack: comprehensive annotation and quantification of small RNA genes. RNA, 19, 740–751.

3. Shahid,S. and Axtell,M.J. (2014) Identification and annotation of small RNA genes using ShortStack. Methods, 67, 20–27.

4. Rogers,K. and Chen,X. (2013) Biogenesis, turnover, and mode of action of plant microRNAs. Plant Cell, 25, 2383–2399.

5. Borges,F. and Martienssen,R.A. (2015) The expanding world of small RNAs in plants. Nat. Rev. Mol. Cell Biol., 16, 727–741.

6. Wierzbicki,A.T., Ream,T.S., Haag,J.R. and Pikaard,C.S. (2009) RNA polymerase V transcription guides ARGONAUTE4 to chromatin. Nat. Genet., 41, 630–634.

7. Liu,Q., Wang,F. and Axtell,M.J. (2014) Analysis of complementarity requirements for plant microRNA targeting using a Nicotiana benthamiana quantitative transient assay. Plant Cell, 26, 741–753.

8. Wang,F., Polydore,S. and Axtell,M.J. (2015) More than meets the eye? Factors that affect target selection by plant miRNAs and heterochromatic siRNAs. Curr. Opin. Plant Biol., 27, 118–124.

9. Zhao,Y., Mo,B. and Chen,X. (2012) Mechanisms that impact microRNA stability in plants. RNA Biol, 9, 1218–1223.

10. Yu,B., Yang,Z., Li,J., Minakhina,S., Yang,M., Padgett,R.W., Steward,R. and Chen,X. (2005) Methylation as a crucial step in plant microRNA biogenesis. Science, 307, 932–935.

11. Li,J., Yang,Z., Yu,B., Liu,J. and Chen,X. (2005) Methylation protects miRNAs and siRNAs from a 3'-end uridylation activity in Arabidopsis. Curr. Biol., 15, 1501–1507.

12. Ren,G., Chen,X. and Yu,B. (2012) Uridylation of miRNAs by hen1 suppressor1 in Arabidopsis. Curr. Biol., 22, 695–700.

13. Zhao,Y., Yu,Y., Zhai,J., Ramachandran,V., Dinh,T.T., Meyers,B.C., Mo,B. and Chen,X. (2012) The Arabidopsis nucleotidyl transferase HESO1 uridylates unmethylated small RNAs to trigger their degradation. Curr. Biol., 22, 689–694.

14. Tu,B., Liu,L., Xu,C., Zhai,J., Li,S., Lopez,M.A., Zhao,Y., Yu,Y., Ramachandran,V., Ren,G., et al. (2015) Distinct and cooperative activities of HESO1 and URT1 nucleotidyl transferases in microRNA turnover in Arabidopsis. PLoS Genet., 11, e1005119.

15. Wang,X., Zhang,S., Dou,Y., Zhang,C., Chen,X., Yu,B. and Ren,G. (2015) Synergistic and independent actions of multiple terminal nucleotidyl transferases in the 3' tailing of small RNAs in Arabidopsis. PLoS Genet., 11, e1005091.

16. Zhai,J., Zhao,Y., Simon,S.A., Huang,S., Petsch,K., Arikit,S., Pillay,M., Ji,L., Xie,M., Cao,X., et al. (2013) Plant microRNAs display differential 3' truncation and tailing modifications that are ARGONAUTE1 dependent and conserved across species. Plant Cell, 25, 2417–2428.

17. Lu,S., Sun,Y.-H. and Chiang,V.L. (2009) Adenylation of plant miRNAs. Nucleic Acids Res., 37, 1878–1885.

18. Patel,P., Ramachandruni,S.D., Kakrana,A., Nakano,M. and Meyers,B.C. (2016) miTRATA: a web-based tool for microRNA Truncation and Tailing Analysis. Bioinformatics, 32, 450–452.

19. Martin,M. (2011) Cutadapt removes adapter sequences from high-throughput sequencing reads. EMBnet.journal, 17, 10–12.

20. Langmead,B., Trapnell,C., Pop,M. and Salzberg,S.L. (2009) Ultrafast and memory-efficient alignment of short DNA sequences to the human genome. Genome Biol., 10, R25.

21. Kozomara,A. and Griffiths-Jones,S. (2014) miRBase: annotating high confidence microRNAs using deep sequencing data. Nucleic Acids Res., 42, D68–73.

22. Blevins,T., Podicheti,R., Mishra,V., Marasco,M., Tang,H. and Pikaard,C.S. (2015) Identification of Pol IV and RDR2-dependent precursors of 24 nt siRNAs guiding de novo DNA methylation in Arabidopsis. Elife, 4, e09591.

23. Zhai,J., Bischof,S., Wang,H., Feng,S., Lee,T.-F., Teng,C., Chen,X., Park,S.Y., Liu,L., Gallego-Bartolome,J., et al. (2015) A One Precursor One siRNA Model for Pol IV-Dependent siRNA Biogenesis. Cell, 163, 445–455.

24. Li,H., Handsaker,B., Wysoker,A., Fennell,T., Ruan,J., Homer,N., Marth,G., Abecasis,G., Durbin,R.1000 Genome Project Data Processing Subgroup (2009) The Sequence Alignment/Map format and SAMtools. Bioinformatics, 25, 2078–2079.

25. Quinlan,A.R. and Hall,I.M. (2010) BEDTools: a flexible suite of utilities for comparing genomic features. Bioinformatics, 26, 841–842.

26. Camacho,C., Coulouris,G., Avagyan,V., Ma,N., Papadopoulos,J., Bealer,K. and Madden,T.L. (2009) BLAST+: architecture and applications. BMC Bioinformatics, 10, 421.

27. Crooks,G.E., Hon,G., Chandonia,J.-M. and Brenner,S.E. (2004) WebLogo: a sequence logo generator. Genome Res., 14, 1188–1190.

28. Stroud,H., Ding,B., Simon,S.A., Feng,S., Bellizzi,M., Pellegrini,M., Wang,G.-L., Meyers,B.C. and Jacobsen,S.E. (2013) Plants regenerated from tissue culture contain stable epigenome changes in rice. Elife, 2, e00354.

29. Shivaprasad,P.V., Chen,H.-M., Patel,K., Bond,D.M., Santos,B.A.C.M. and Baulcombe,D.C. (2012) A microRNA superfamily regulates nucleotide binding siteleucine-rich repeats and other mRNAs. Plant Cell, 24, 859–874.

30. Shivaprasad,P.V., Dunn,R.M., Santos,B.A., Bassett,A. and Baulcombe,D.C. (2012) Extraordinary transgressive phenotypes of hybrid tomato are influenced by epigenetics and small silencing RNAs. EMBO J, 31, 257–266.

31. Fei,Q., Li,P., Teng,C. and Meyers,B.C. (2015) Secondary siRNAs from Medicago NB-LRRs modulated via miRNA-target interactions and their abundances. Plant J, 83, 451–465.

32. Coruh,C., Cho,S.H., Shahid,S., Liu,Q., Wierzbicki,A. and Axtell,M.J. (2015) Comprehensive Annotation of Physcomitrella patens Small RNA Loci Reveals That the Heterochromatic Short Interfering RNA Pathway Is Largely Conserved in Land Plants. Plant Cell, 27, 2148–2162.

33. Xie,Z.X., Johansen,L.K., Gustafson,A.M., Kasschau,K.D., Lellis,A.D., Zilberman,D., Jacobsen,S.E. and Carrington,J.C. (2004) Genetic and functional diversification of small RNA pathways in plants. Plos Biology, 2, 642–652.

34. Henderson,I.R., Zhang,X., Lu,C., Johnson,L., Meyers,B.C., Green,P.J. and Jacobsen,S.E. (2006) Dissecting Arabidopsis thaliana DICER function in small RNA processing, gene silencing and DNA methylation patterning. Nat. Genet., 38, 721–725.

35. Wang,H., Zhang,X., Liu,J., Kiba,T., Woo,J., Ojo,T., Hafner,M., Tuschl,T., Chua,N.-H. and Wang,X.-J. (2011) Deep sequencing of small RNAs specifically associated with Arabidopsis AGO1 and AGO4 uncovers new AGO functions. Plant J, 67, 292–304.

36. Mi,S., Cai,T., Hu,Y., Chen,Y., Hodges,E., Ni,F., Wu,L., Li,S., Zhou,H., Long,C., et al. (2008) Sorting of small RNAs into Arabidopsis argonaute complexes is directed by the 5' terminal nucleotide. Cell, 133, 116–127.

37. Li,S., Vandivier,L.E., Tu,B., Gao,L., Won,S.Y., Li,S., Zheng,B., Gregory,B.D. and Chen,X. (2015) Detection of Pol IV/RDR2-dependent transcripts at the genomic scale in Arabidopsis reveals features and regulation of siRNA biogenesis. Genome Res., 25, 235–245.

38. Yang,D.-L., Zhang,G., Tang,K., Li,J., Yang,L., Huang,H., Zhang,H. and Zhu,J.-K. (2016) Dicer-independent RNA-directed DNA methylation in Arabidopsis. Cell Res., 26, 66–82.

39. Ye,R., Chen,Z., Lian,B., Rowley,M.J., Xia,N., Chai,J., Li,Y., He,X.-J., Wierzbicki,A.T. and Qi,Y. (2016) A Dicer-Independent Route for Biogenesis of siRNAs that Direct DNA Methylation in Arabidopsis. Mol. Cell, 61, 222–235.

40. Cho,S.H., Addo-Quaye,C., Coruh,C., Arif,M.A., Ma,Z., Frank,W. and Axtell,M.J. (2008) Physcomitrella patens DCL3 is required for 22-24 nt siRNA accumulation, suppression of retrotransposon-derived transcripts, and normal development. PLoS Genet., 4, e1000314.

